# Coupling adipose tissue architecture and metabolism via cytoophidia

**DOI:** 10.1101/2021.12.02.470854

**Authors:** Jingnan Liu, Yuanbing Zhang, Youfang Zhou, Qiao-Qi Wang, Kang Ding, Suwen Zhao, Pengfei Lu, Ji-Long Liu

**Author notes:** These authors contributed to this work equally (Jingnan Liu, Yuanbing Zhang).

## Abstract

Tissue architecture determines its unique physiology and function. How these properties are intertwined has remained unclear. Here, we show that the metabolic enzyme CTP synthase (CTPS) form filamentous structures termed cytoophidia along the adipocyte cortex in *Drosophila* adipose tissue. Interestingly, loss of cytoophidia, whether due to reduced CTPS expression or a point mutation that specifically abrogates its polymerization ability, leads to downregulated Collagen-Integrin signaling, weakened adipocyte adhesion, and defective adipose architecture. Strikingly, CTPS specifically binds with Integrin subunit α2, which influences Integrin function and Collagen IV deposition. cytoophidia promote Collagen IV mRNA expression and thus its extracellular deposition to strengthen adipocyte adhesion. Remarkably, Collagen IV-Integrin signaling reciprocally regulates cytoophidium formation at a post-translational level. Together, we demonstrate that a positive feedback signaling loop containing both cytoophidia and Integrin adhesion complex couples tissue architecture and metabolism in the fly adipose.

## INTRODUCTION

Tissue structure or architecture refers to the organization of individual cells and their extracellular matrix (ECM) in such a way that it is best suited to perform the unique physiological function of each tissue. It is mainly accomplished, irrespective of whether the tissue of concern is an epithelium or a connective tissue such as the adipose, by tissue-specific regulation of cell-cell and cell-ECM adhesion (Mian et al. 2003; Pozzi et al. 2017). Despite its importance, how tissue architecture determines or is determined by its physiological functions, for example, its metabolism, has remained largely unclear.

As a metabolic organ and an equivalent of the mammalian adipose tissue, the *Drosophila* fat body has in recent years emerged as an excellent model to understand fat metabolism, due in part to its relatively simple and well-defined tissue organization and architecture (Baker and Thummel 2007; Arrese and Soulages 2010; Li et al. 2019). Briefly, the fly fat body is a single-layered tissue of adipocytes encapsulated by a dense basement membrane (BM) (Baker and Thummel 2007; Pope et al. 2016). Similar to the BM anchoring the basal side of an epithelium, the BM in the fat is also composed of multiple ECM components, including the Laminin and the Collagen IV network of polymers, which are interconnected by numerous “adaptor/linker” proteins such as Nidogen and Perlecan (Cummings et al. 2016; Jayadev and Sherwood 2017). *Drosophila* Collagen IV polymers are formed by an α1 chain and an α2 chain (Natzle et al. 1982; Fessler and Fessler 1989). In addition to contributing to forming the BM, Collagen IV and other ECM components are also deposited between adipocytes. Indeed, intercellular deposition of Collagen IV, via binding to its membrane-receptor Integrin, is essential for cell-matrix adhesion and the fly adipose tissue organization (Dai et al. 2017).

As the primary ECM receptor, Integrins are heterodimeric membrane proteins composed of an α and β subunit (Shattil et al. 2010). In *Drosophila*, the Integrin family consists of five αPS subunits (αPS1 to 5) and two β subunits (βPS and βν) (Brown 1993; Brower et al. 1995). Together with its co-receptor Syndecan (Sdc) and cytoplasmic downstream components, including Integrin-linked <u>k</u>inase (ILK), <u>P</u>articularly <u>in</u>teresting <u>c</u>ysteine-<u>h</u>istidine-rich protein (PINCH), and Talin, Integrins form an “adhesion complex,” which binds to the actin cytoskeleton and transduces environmental signals to the cytoplasm and the nucleus (Hannigan et al. 2005; Morgan et al. 2007; Dai et al. 2017). This integrin adhesion complex thus serves as a gateway linking both the microenvironment and the inside of a cell and plays an essential role in mechanosensing, cell migration, differentiation, etc. (Diamond and Springer 1994; Giancotti and Ruoslahti 1999; Hynes 2002; Campbell and Humphries 2011; Kechagia et al. 2019; Moreno-Layseca et al. 2019).

The metabolic enzyme Cytidine 5’-triphosphate synthase (CTPS) catalyzes the conversion of UTP to CTP, a rate-limiting step in the *de novo* synthesis of CTP (Lieberman 1956; Levitzki and Koshland 1969). The active enzyme is a homo-tetramer with each monomer consisting of a glutamine aminotransferase domain, a kinase-like ammonia ligase domain, and an α-helical linker (Levitzki and Koshland 1976; Robertson 1995). Remarkably, it has been widely observed in both prokaryotes and eukaryotes that CTPS often forms filamentous structures termed cytoophidia although the functional significance of filament formation has remained unknown (Ingerson-Mahar et al. 2010; Liu 2010; Noree et al. 2010; Carcamo et al. 2011; Liu 2011; Liu 2016; Daumann et al. 2018; Zhou et al. 2020).

We recently found that cytoophidia exist in the fly fat body (Zhang et al. 2020). Here, we hypothesized that cytoophidia are essential for the fly adipocytes. Surprisingly, we found that cytoophidia regulate Integrin and Collagen IV function and play an important role in fat body adipocyte adhesion and tissue architecture.

## RESULTS

### Cytoophidia localize at the cortex of fly adipocytes

We first determined CTPS expression in the fat body during various stages of fly development. To visualize the subcellular localization and dynamics of endogenous CTPS *in vivo*, we created a “knock-in” fly line in which the coding sequences of the fluorescent protein mCherry and the V5 tag were inserted in-frame at the C-terminus of CTPS, referred hereafter to the *CTPS-* mCh line. Using Western blotting analysis, we confirmed that the V5-tagged CTPS protein was expressed at the first-instar larval (L1), the second-instar larval (L2), and the early, middle, and late third-instar larval (L3) stages (Fig. 1A, B). Using fluorescent confocal microscopy, we found that cytoophidia were present at all of the stages examined (Fig. 1C). Interestingly, however, both the lengths and numbers of cytoophidia progressively reduced from the early L3 stage to the late L3 stage (Fig. 1D, E).

**Figure 1.**
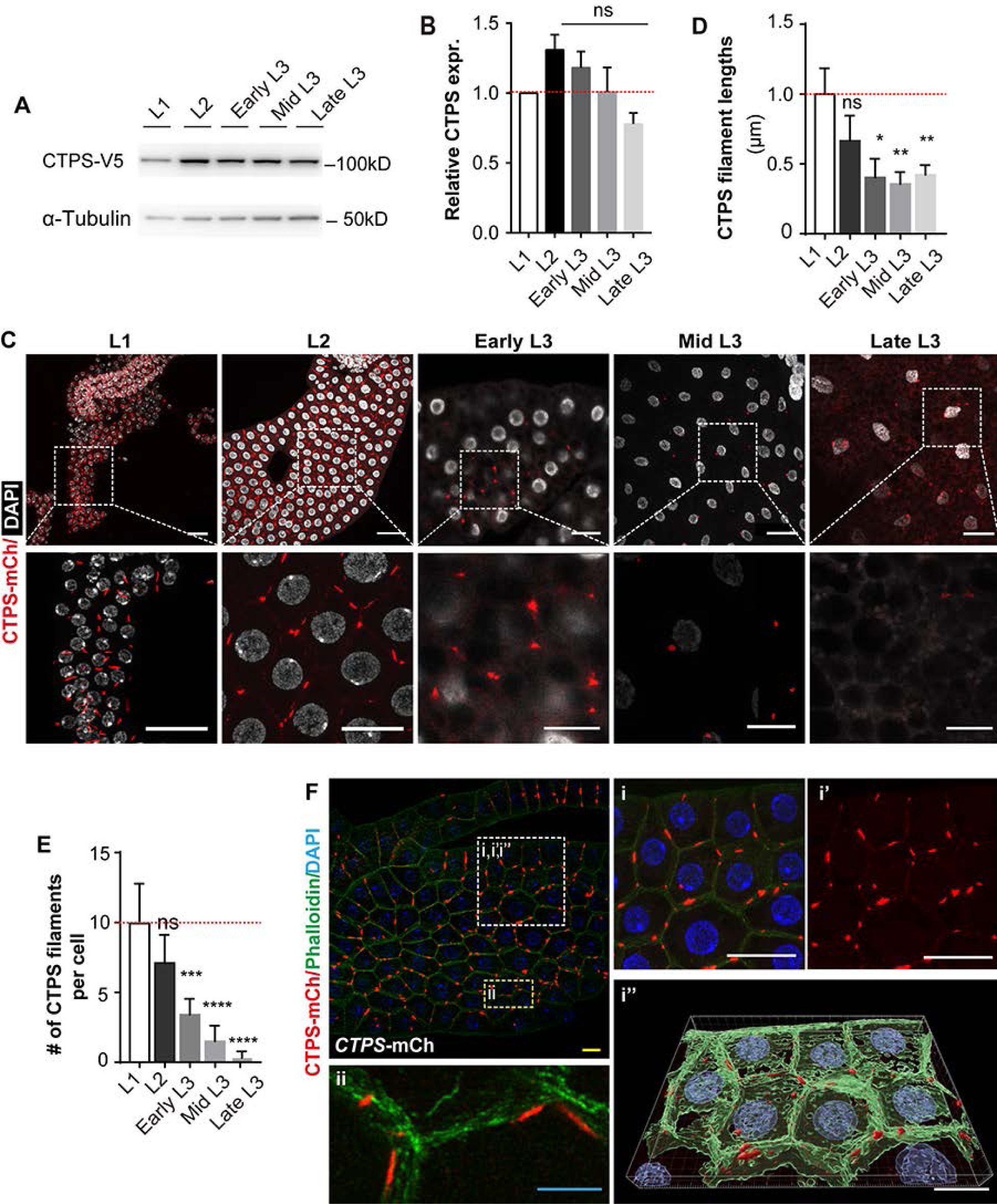
Cytoophidia localize at the cortex of fly adipocytes. (**A-B**) Western blotting analysis of CTPS proteins from fat bodies of *CTPS*-mCh larvae (**A**, 20 flies/genotype). (**B**) Quantification of CTPS relative expression levels at various stages when compared with that at the L1 stage. Anti-V5 antibody was used for the immunoblotting analysis. Alpha-tubulin was used as an internal control. (**C**) Confocal images of the fat bodies from the first-, second- and third-instar larvae. Early L3, 72 hours after egg laying (AEL); Mid L3, 96 hours AEL; Late L3, 120 hours AEL. Nuclei were stained with DAPI (white). (n = 20 flies/genotype, representative of three independent experiments). Areas in green dotted boxes were zoomed into for closers views of cytoophidia (red) and the nuclei (white). Scale bars, 20 μm. (**D, E**) Quantification of filament lengths (**D**) and numbers (**E**) per adipocytes at the above stages. Values were normalized with those from the L1 stage. (**F**) Confocal images of a fat body from a *CTPS*-mCh larva at the second-instar stage. Note that actin cytoskeleton, marked by phalloidin staining, was enriched at cell cortex. Nuclei were stained with DAPI (blue). The area in a white dotted square is magnified in closed-up views on the right panels (i and i’), and processed by Imaris (ii’). The area in a yellow dotted square is viewed at a high resolution using STED microscopy in the lower left panel (ii). Scale bars, 5μm (blue), 20 μm (white) and 50 μm (yellow). All values are the means ±S.E.M. ns, no significance, *p<0.05, **p<0.01, *** p < 0.001, **** p < 0.0001 by Student’s *t*-test.

Next, we used Stimulated Emission Depletion (STED) microscopy to examine the subcellular localization of cytoophidia at a high resolution. We found that almost all cytoophidia were located at the cell cortex, as marked by intense phalloidin staining of the actin filaments (Fig. 1Fi-ii). This was in contrast to the nuclei, which were centrally located in the adipocytes (Fig. 1Fi-ii). To determine whether this striking feature of cytoophidium localization was an artifact introduced by protein fusion between CTPS and mCherry or the V5 tag, we performed immunofluorescence microscopy and directly detected CTPS protein expression in the fat body of the wild type *w^1118^* fly line at the L2 stage. Again, we found that cytoophidia, as detected by the CTPS antibody, were present at the adipocyte cortex (Supplemental Fig. 1A), thus confirming the results from the *CTPS-*mCh knock-in line. Together, the results show that cytoophidia are present in the fly fat-body adipocytes and that they align along the cortex of adipocytes.

### CTPS reduction causes defective adipocyte adhesion

The cortical localization of cytoophidia correlated with adipocyte adhesion sites. Therefore, we asked whether cytoophidia play a role in adipocyte adhesion. To answer this question, we used the RNAi technology and created a fly line in which CTPS expression levels were specifically reduced in the fat body under the control of the *Cg GAL4* driver (*Cg GAL4*>*CTPS*-RNAi), referred to as the *CTPS-*Ri hereafter. Using qPCR, we found that *CTPS* mRNA expression in the fat body was reduced by ∼73% in the *CTPS-* Ri larvae when compared with the wild-type larvae (*Cg GAL4*>*w^1118^*) at the L3 stage (Fig. 2A). Unlike the control fat body at this stage, which was a thin, tight, ribbon-like structure, we found that the *CTPS-*Ri fat body was loosely connected, often with adipocytes breaking off of the adipose tissue (Fig. 2B). We speculated that the mutant fat-body suffered from an adhesion defect where adipocytes were not as tightly connected as they should.

**Figure 2.**
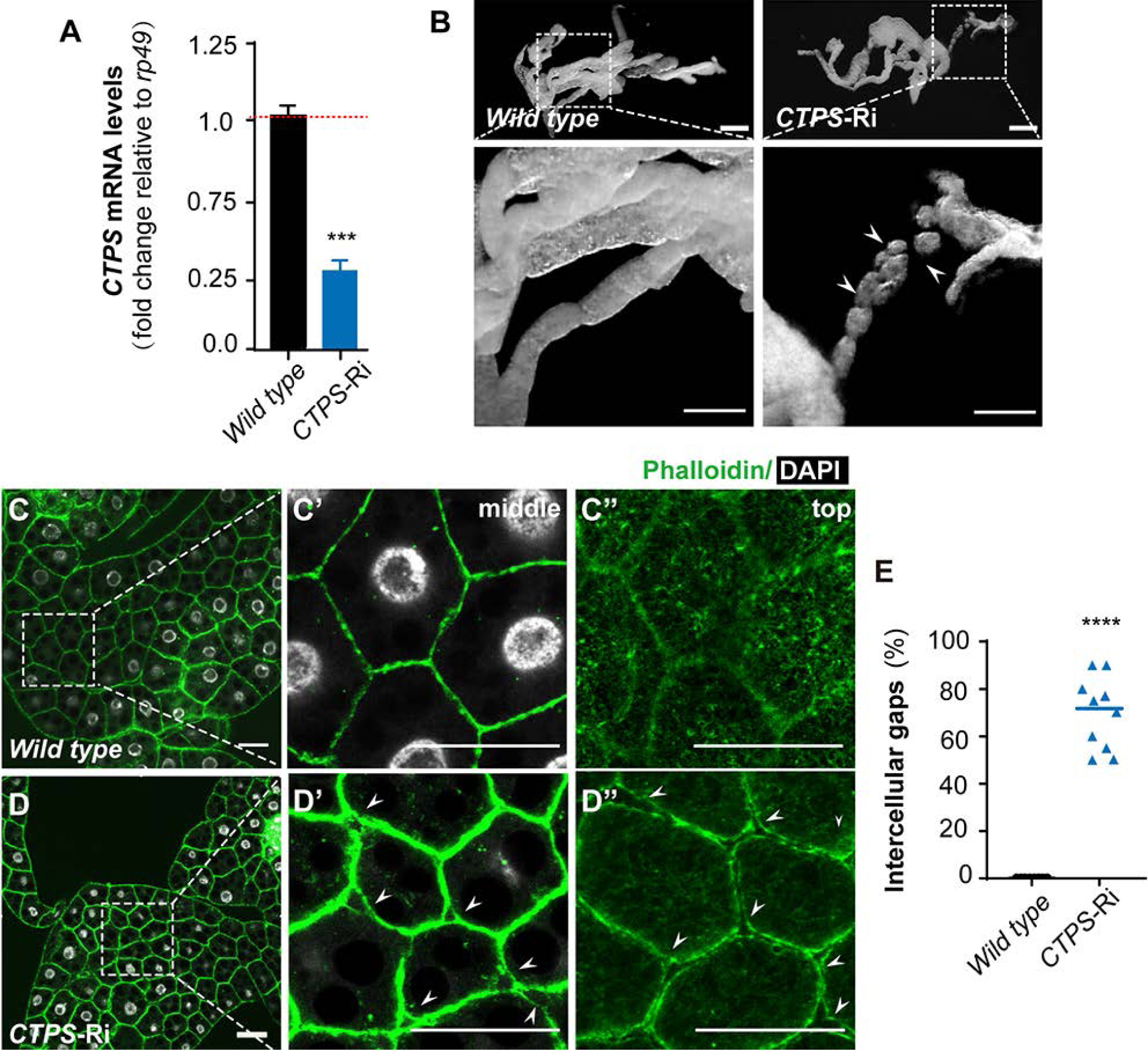
CTPS reduction causes defective adipocyte adhesion. **(A)** Relative *CTPS* mRNA expression levels as measured by quantitative RT-PCR using fat bodies from the wild-type (*CgG4*>*w^1118^*) and *CTPS*-Ri larvae at the third instar stages (10 flies/genotype; n=3). **(B)** Wholemount images of the fat bodies from the wild-type and (*CgG4*>*CTPS-*Ri) larvae at the third instar stage. Areas in the yellow dotted boxes were shown in close-up views to show more clearly tissue morphologies. Note that disjointed fat body tissues from the mutant larvae, indicated weakened tissue connection. Scale bars, 250 μm. (**C**-**E**) Adipose tissue morphology as examined by fluorescent confocal microscopy using the wild-type (**C**) and *CTPS*-Ri fat bodies (**D**) from the third instar stage. Phalloidin and DAPI were used to stain actin cytoskeleton and nuclei, respectively. Note that, while wild-type adipocytes were compact and tightly joined, *CTPS*-Ri adipocytes were loosely connected with numerous gaps among them at both the middle (**C’**, **D’**) and the top (**C’’**, **D’’**) focal planes along the Z-axis (perpendicular to the single-layered fat-body). Intercellular gaps quantified in (**E**), indicate the percentage of adipocytes that have gaps. Each point represents the number of gaps in a single fat body image. All values are the means ±S.E.M. *** p < 0.001, **** p < 0.0001 by Student’s t-test.

Therefore, we next examined the tissue integrity of the *CTPS-*Ri fat body at the L2 stage, before the tissue dissociation phenotype became apparent at the L3 stage. Using fluorescent confocal microscopy to examine the actin cytoskeleton and the nuclei, we observed the typical polygonal adipocytes with tight cell-cell contacts in the wild-type *w^1118^* fat body (Fig. 2C). Remarkably, the mutant fat body contained many gaps at the bi-cellular or tri-cellular contact sites (Fig. 2D). By performing Z-stack image acquisition and reconstruction, we confirmed that intercellular gaps were present throughout the Z-axis, including in the middle and at the top of the single-cell-layered adipose tissue (Fig. 2D, E). Together, our results show that CTPS reduction leads to weakened adipocyte adhesion and dissociation of the adipose tissue in the fat body.

### CTPS polymerization promotes its protein levels

Next, we asked whether it is the filaments, rather than the protein expression level *per se*, of CTPS that are essential for adipocyte adhesion. To answer this question, we needed to specifically disrupt the polymerization of CTPS proteins without affecting their enzymatic functions. Previous studies showed that the Histidine amino acid at the 355th position, or His^355^, of human CTPS protein is essential for its polymerization, but not enzymatic function (Lynch et al. 2017; Sun and Liu 2019). By aligning the amino-acid sequences of both human and fly CTPS proteins and performing structural analysis, we identified that the His^355^ amino acid of the fly CTPS in the glutamine amidotransferase domain would also be at the interface between two consecutive tetramers and thus should also be critical for filament formation (Fig. 3A, B).

**Figure 3.**
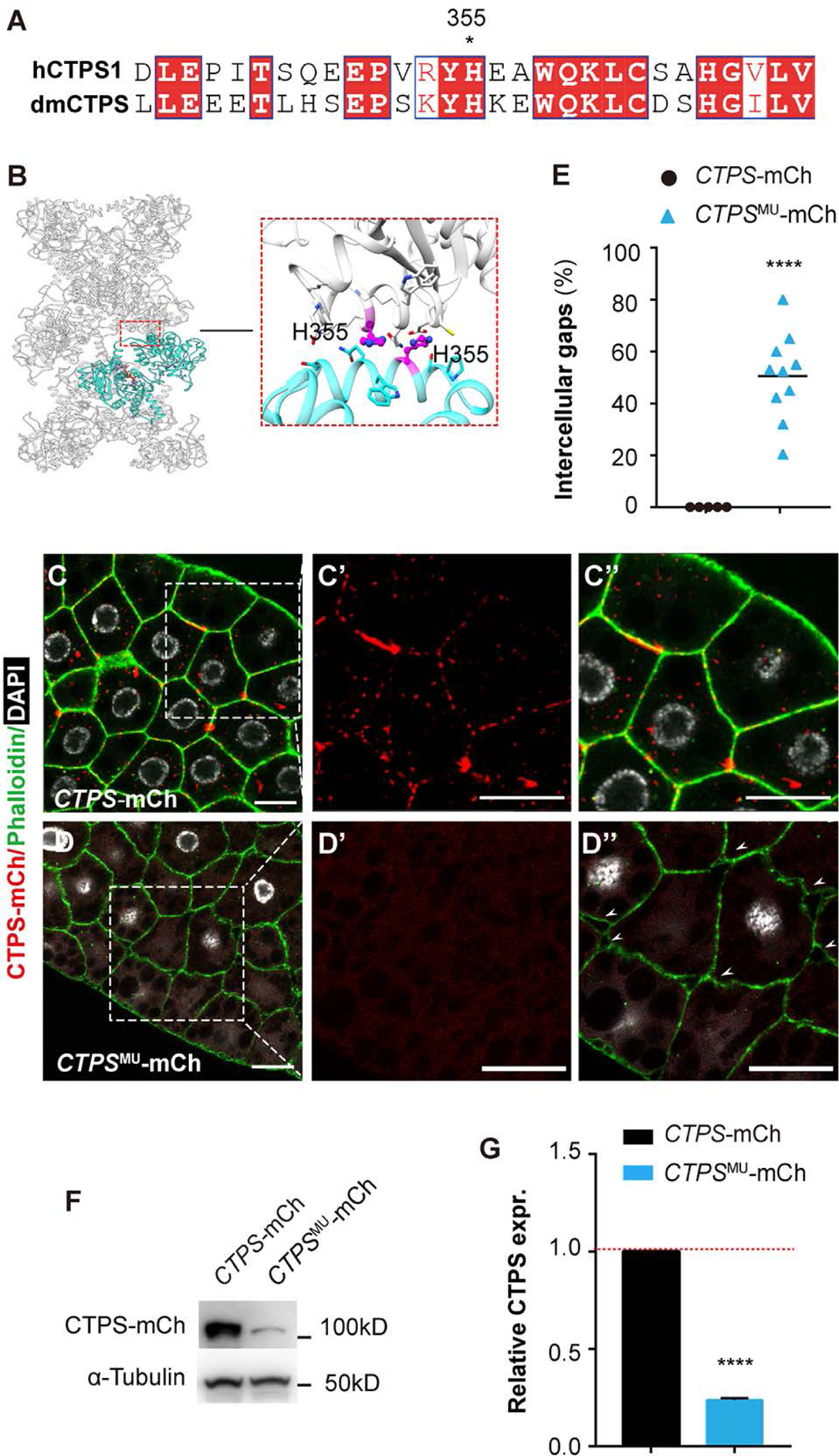
CTPS polymerization promotes its protein levels. (**A**) Alignment of amino acid sequences of *Drosophila* and Human CTPS proteins indicating the conserved Histidine at the 355 position. (**B**) Structural modeling of CTPS protein indicates that H^355^ locates at the interface of two consecutive tetramers in the glutamine aminotransferase domain. (**C**-**E**) Adipose tissue morphology as examined by fluorescent confocal microscopy using the control (*CTPS*-mCh) (**C**) and the mutant (*CTPS*^MU^-mCh) fat bodies (**D**) from the third instar stage. Phalloidin and DAPI were used to stain actin cytoskeleton and nuclei, respectively. Note the prominent presence of cytoophidia (red) and tight adipocyte adhesion in the control (**C**-**C**’) and the absence of cytoophidium and presence of numerous gaps in the mutant fat bodies (**C-D’**). Intercellular gaps quantified in (**E**), indicate the percentage of adipocytes that have gaps. Each point represents the number of gaps in a single fat body image. (**F**) Immunoblotting analysis of CTPS proteins from larval lysate at the early 3rd instar stage (30 flies/genotype, n=3). Anti-mCherry antibody was used to detect the expression of CTPS-mCherry fusion protein. Alpha-tubulin was used as an internal control. (**G**) Quantification of the relative expression levels of CTPS protein. All values are the means ±S.E.M. **** p < 0.0001 by Student’s t-test.

We predicted that a replacement of His^355^ with Alanine, which is uncharged and smaller than Histidine, would disrupt the polymerization ability of the fly CTPS protein in a similar way to the human CTPS situation. Thus, using the CRISPR/Cas9-mediated knock-in technology, we created a point mutant that would lead to His^355^ to Ala^355^ conversion in the endogenous CTPS protein. Moreover, the mutant CTPS was fused with mCherry and three hemolymph agglutinin tags (3x HA) at its C-terminus so that the fusion protein could be easily detected in various assays (Supplemental Fig. 2). Using the genomic

DNA extracted from the knock-in fly line, referred to as the *CTPS*^MU^-mCh line hereafter, we confirmed the intended point mutation was successfully introduced as designed (Supplemental Fig. 3). The homozygous *CTPS*^MU^-mCh adult flies were viable, with no discernible developmental defects other than a smaller body size.

Next, we examined the adipose tissues in both the control *CTPS-*mCh and the *CTPS*^MU^-mCh flies. As expected, we observed numerous cytoophidia in the control adipocytes, but none in *CTPS*^MU^-mCh adipocytes (Fig. 3C, C’, D, D’). These results thus confirmed our above prediction that the mutant CTPS proteins would be unable to polymerize, and suggest that, as in the human situation, His^355^ is also essential for fly CTPS proteins to form filaments.

Furthermore, phalloidin staining of the actin cytoskeleton showed that numerous gaps emerged between adipocytes in the mutant but not the control adipose tissues, indicating that adipocyte adhesion is defective in the mutant fly line (Fig. 3C’-E). Interestingly, CTPS protein levels, as detected by Western blotting analysis using an antibody against the mCherry fusion protein, were greatly reduced in the *CTPS*^MU^-mCh fly larvae when compared with the control (Fig. 3F, G). The data thus show that a failure to form CTPS polymers leads to a reduction in its protein levels.

Thus, while adipocyte adhesion is defective in the fat body of the *CTPS*^MU^-mCh fly larvae, we concluded that the defect is most likely a secondary consequence caused by reduced CTPS expression, rather than a direct result from a lack of cytoophidia in the mutant flies. Moreover, the data suggest that filament formation increases CTPS protein levels in fly adipocytes, presumably by stabilizing the otherwise labile unpolymerized proteins.

### Cytoophidia, rather than CTPS protein levels, are essential for adipocyte adhesion

To definitively determine whether cytoophidia are essential for adipocyte adhesion, we needed to disrupt filament formation without reducing its protein expression levels. To this end, we created a transgenic fly line, the *CgG4*/CTPS^MU-OE^, in which a mutant CTPS protein, tagged with mCherry-HA at its C-terminus, was forcefully expressed in the fat body as controlled by the *Cg GAL4* driver. We used Arg^355^, which is negatively charged and is bulkier than Alanine used above, to replace His^355^ to ensure that the mutant CTPS protein is incapable of forming filaments. As a control, a transgenic line *CgG4*/CTPS^WT-OE^, in which the wild-type CTPS tagged with mCherry-HA was also created. Western blotting analysis confirmed that protein expression of the mutant CTPS was at a similar level as the wild-type CTPS (Fig. 4A, B).

**Figure 4.**
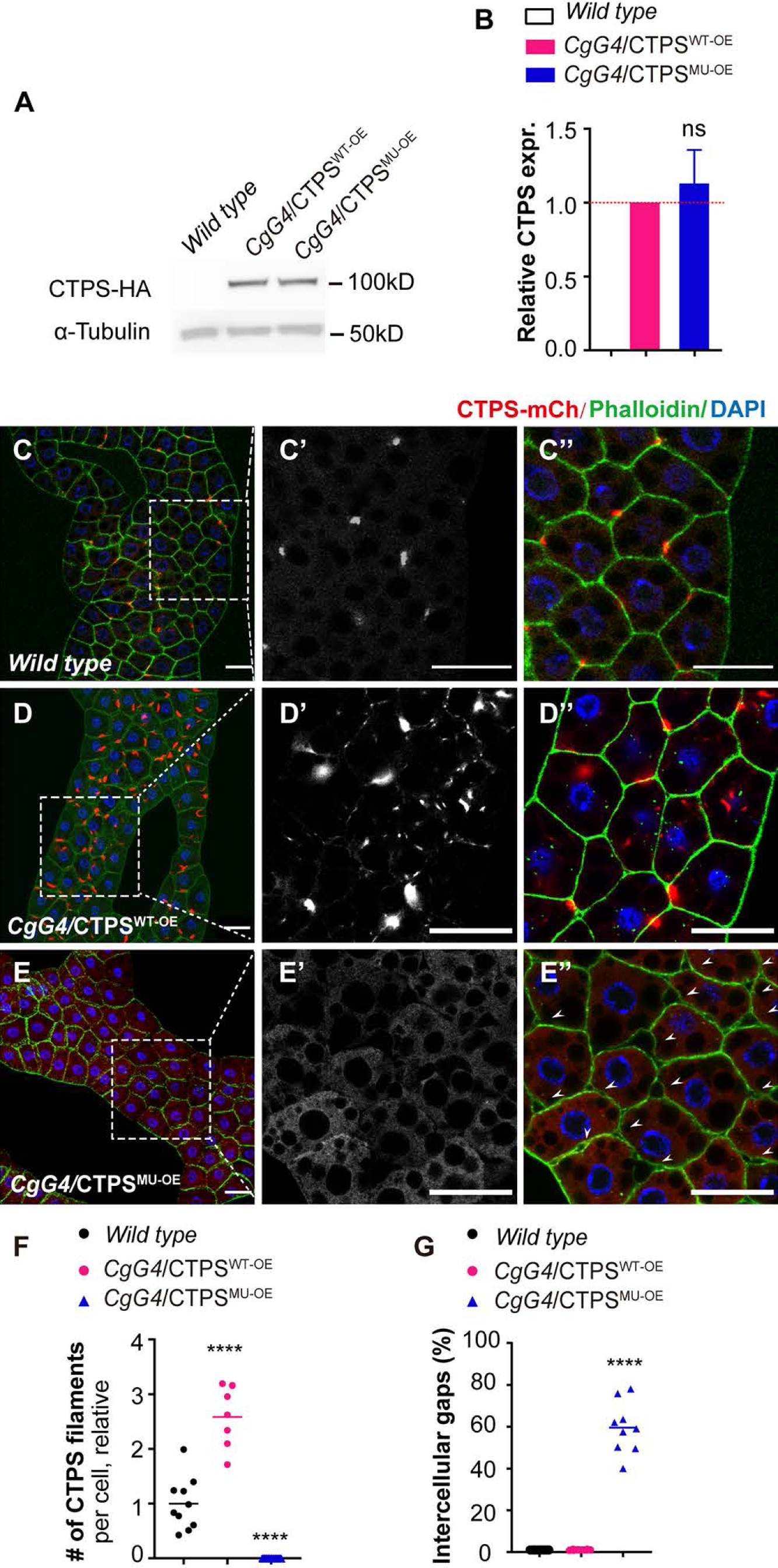
Cytoophidia, rather than CTPS protein levels, are essential for adipocyte adhesion. (**A**) Immunoblotting analysis of CTPS proteins from larval lysate at the early 3rd instar stage (30 flies/genotype, n=2). Anti-HA antibody was used to detect the expression of CTPS-mCherry fusion protein. Alpha-tubulin was used as an internal control. (**B**) Quantification of the relative expression levels of CTPS protein. (**C**-**E**) Fluorescent confocal images of the fat bodies from the wild-type (*w^1118^*) (**C**-**C’’**), the CTPS^WT-OE^ (**D**-**D’’**), and the CTPS^MU-OE^ (**E**-**E’’**) fly lines carrying the *CTPS*-mCh allele at the third instar stage. Phalloidin and DAPI were used to stain actin cytoskeleton and nuclei, respectively. Note the prominent presence of cytoophidia (red) in the wild-type (**C’’**) and CTPS^WT-OE^ (**D’’**) fat bodies, with the latter having a more filaments than the former. Also note that cytoophidia were absent from the CTPS^MU-OE^ line (**E’’**) and adipocyte adhesion was defective. White arrows indicate intercellular gaps. Scale bars, 20 μm. (**F**, **G**) Quantification of the numbers of filaments (**F**) or intercellular gas (**G**) per adipocyte. All values are the means ±S.E.M. ns, no significance, **** p < 0.0001 by Student’s t-test.

As expected, cytoophidia were present in the adipocytes of both the *CTPS-* mCh and the *CgG4/CTPS*^WT-OE^ fly lines, with the latter having a larger than normal number of filaments (Fig. 4C-D’’, F). There were no gaps between adipocytes, and adipose morphology was normal in *CgG4/*CTPS^WT-OE^ fly lines, as did wild-type adipocytes of the *w^1118^* line (Fig. 4C-D’’, G). By contrast, no filaments at all were observed in the adipocytes of the *CgG4*/CTPS^MU-OE^ adipose tissue (Fig. 4E-E’, F), confirming that His^355^ to Arg^355^ conversion abrogates the ability of CTPS to form filaments in fly adipocytes. Importantly, we observed gaps between adipocytes in the *CgG4*/CTPS^MU-OE^ adipose tissue (Fig. 4E’’, G) similar to what we described in the flies with a reduction of CTPS proteins (Fig. 2D’’, E). Together, the data demonstrate that cytoophidia, rather than the absolute amount of CTPS proteins, are essential for adipocyte adhesion.

### Cytoophidia promote Collagen IV production by up-regulating its mRNA expression

Next, we sought to determine whether cytoophidia promote adipocyte adhesion by regulating Collagen IV-Integrin signaling. To this end, we first examined whether cytoophidia affected Integrin expression. To track Integrin expression, we used the *If*-GFP fluorescent trap line, which marks the integrin αPS2 subunit Inflated (If) protein, and crossed it with fly lines carrying either the *CTPS-*mCh, the *CTPS*^MU^-mCh, the *CgG4/*CTPS^WT-OE^, or the *CgG4/*CTPS^MU-OE^ allele. Integrin expression, as marked by GFP, was found at the cell periphery, presumably the plasma membrane, of all four fly lines (Fig. 5A’-D’). Interestingly, the levels of Integrin protein expression, as indicated by If-GFP fluorescent intensity, were not significantly changed in the *CTPS*^MU^-mCh or the *CgG4/*CTPS^MU-OE^ adipocytes, except in the *CgG4/*CTPS^WT-OE^ adipocytes where it is significantly increased, when compared to the control (Fig. 5A’-D’, I).

**Figure 5.**
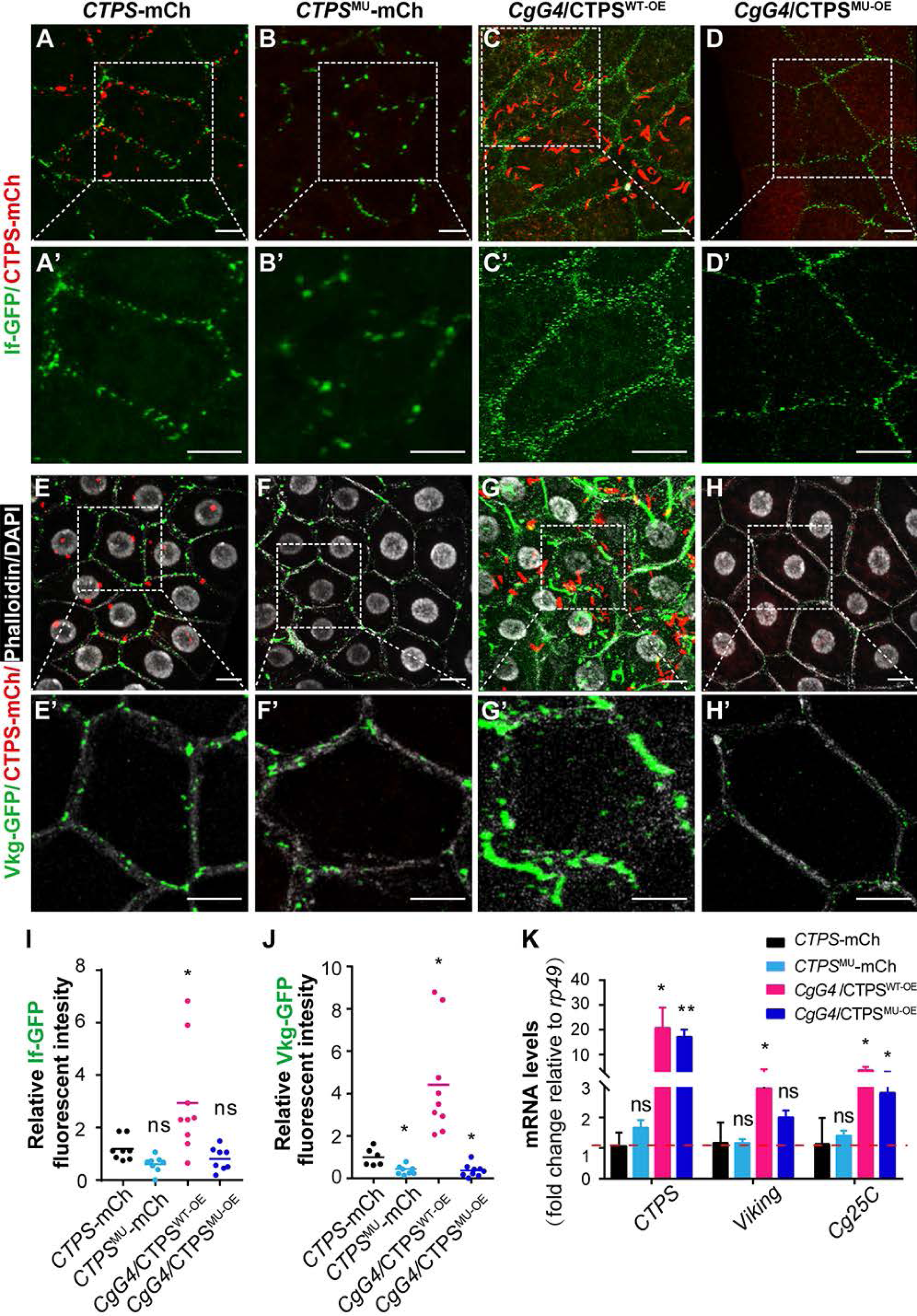
Cytoophidia promote Collagen IV production by up-regulating its mRNA expression. (**A**-**D’**) Fluorescent confocal images showing Integrin expression, as reported by the *If*-GFP trap line, in the fly lines carrying either the *CTPS-*mCh (**A**, **A’**), the *CTPS*^MU^-mCh (**B**, **B’**), the *CgG4/*CTPS^WT-OE^ (**C**, **C’**), or the *CgG4/*CTPS^MU-OE^ allele (**D**, **D’**). Note that Integrin expression was at the cell periphery, presumably the plasma membrane. Areas in white dotted boxes were shown in close-up views to indicate more clearly Integrin expression. (**I**) Relative Integrin expression as measured by fluorescent intensity in the adipocytes of the four fly lines. Scale bars, 25 µm. (**E**-**H’**) Fluorescent confocal images showing Collagen IV expression, as reported by the *Vkg*-GFP trap line, in the fly lines carrying either the *CTPS-* mCh (**E**, **E’**), the *CTPS*^MU^-mCh (**F**, **F’**), the *CgG4/*CTPS^WT-OE^ (**G**, **G’**), or the *CgG4/*CTPS^MU-OE^ allele (**H**, **H’**). Areas in white dotted boxes were shown in close-up views to indicate w more clearly Collagen IV expression. (**J**) Relative Integrin expression as measured by fluorescent intensity in the adipocytes of the four fly lines. Scale bars, 25 µm. (**K**) qPCR analysis of the relative levels of mRNA expression of indicated genes. Samples were fat body lysate from the early third instar larvae of the indicated genotypes. Values were normalized to those from the *CTPS*-mCh line. All values were the means ± S.E.M. ns, no significance, * p < 0.05, ** p < 0.01 by one-way ANOVA with a Tukey post hoc test.

Using a similar strategy, we examined whether cytoophidia affected Collagen IV production. To mark collagen IV expression, we used the *Vkg*-GFP fluorescent trap line and crossed it with flies carrying either the *CTPS-* mCh, the *CTPS*^MU^-mCh, the *CgG4/*CTPS^WT-OE^, or the *CgG4/*CTPS^MU-OE^ allele. As expected, we observed abundant Collagen IV expression in adipocytes of the *CTPS-*mCh larvae (Fig. 5E, E’). Interestingly, however, comparing with the control, Collagen IV expression was reduced in adipocytes of the *CTPS*^MU^-mCh and the *CgG4/*CTPS^MU-OE^ fly lines but was greatly increased in adipocytes of the *CgG4-*CTPS^WT-OE^ fly line (Fig. 5E-H’, J). Thus, the amount of cytoophidia appeared to correlate with Collagen IV levels, suggesting that cytoophidia promote Collagen IV protein expression.

Next, we examined whether cytoophidia regulate the expression of other ECM components, for example, Nidogen. To this end, we crossed the *Ndg*-sGFP fluorescent trap line, in which Nidogen is marked by sGFP expression, with the above control and experimental fly lines. We found that Nidogen expression levels were not significantly changed in adipocytes of the *CTPS*^MU^-mCh, the *CgG4/*CTPS^WT-OE^, or the *CgG4/*CTPS^MU-OE^ flies when compared to the control *CTPS-*mCh flies (Supplemental Fig. 4A-E). These data thus suggest that cytoophidia specifically promote the expression of Collagen IV, but not other ECM components.

To determine whether cytoophidia regulate Collagen IV expression at the transcriptional level, we used qPCR and assessed *Viking* and *Cg25C* mRNA expression in the fat-body adipocytes from the *CTPS-*mCh, the *CTPS*^MU^-mCh, the *CgG4/*CTPS^WT-OE^, and the *CgG4/*CTPS^MU-OE^ fly lines. As a control, we also examined *CTPS* mRNA expression. As expected, we found that *CTPS* mRNA expression was similar in the *CTPS*^MU^-mCh adipocytes when compared to the *CTPS-*mCh adipocytes, but it was greatly upregulated in adipocytes of both the *CgG4/CTPS*^WT-OE^ and the *CgG4/*CTPS^MU-OE^ fly lines (Fig. 5K). Remarkably, *Viking* mRNA expression was greatly upregulated in the *CgG4/*CTPS^WT-OE^ adipocytes, but not significantly changed in either the *CTPS*^MU^-mCh or the *CgG4/*CTPS^MU-OE^ adipocytes when compared to the *CTPS-*mCh adipocytes (Fig. 5K). Similar results were also observed when we examined *Cg25C* mRNA expression, although it was also significantly increased in the *CgG4/*CTPS^MU-OE^ adipocytes (Fig. 5K).

Together, the combination of loss- and gain-of-function studies of CTPS conclusively demonstrate that cytoophidia specifically promote the production of Collagen IV, rather than other ECM components such as Nidogen. Further, the data show that cytoophidia regulate Collagen IV expression at the transcriptional level.

### Integrin signaling promotes cytoophidium formation

The localization of cytoophidia at the adipocyte cortex, where cell-ECM adhesion occurs, prompted us to ask whether Collagen IV-integrin signaling, which mediates adipocyte adhesion, regulates cytoophidium formation. To this end, we crossed the *CTPS-*mCh line with fly lines with reduced mRNA expression via RNAi of either the Integrin αPS2 subunit (If), or the Integrin βPS subunit (myospheroid, Mys), or PINCH (encoded by the steamer duck gene, *Stck*), or ILK. As expected, adipocyte adhesion was defective, as evident from the presence of intercellular gaps, in fly lines carrying any of the RNAi alleles, including the *If-Ri*, *Mys-Ri*, *Stck-Ri*, and *Ilk-Ri* alleles when compared with the wild-type control (Fig. 6A-E, K). Remarkably, cytoophidia were absent in the adipocytes in all of these transgenic fly lines with reduced integrin signaling as well (Fig. 6A’-E’, L).

**Figure 6.**
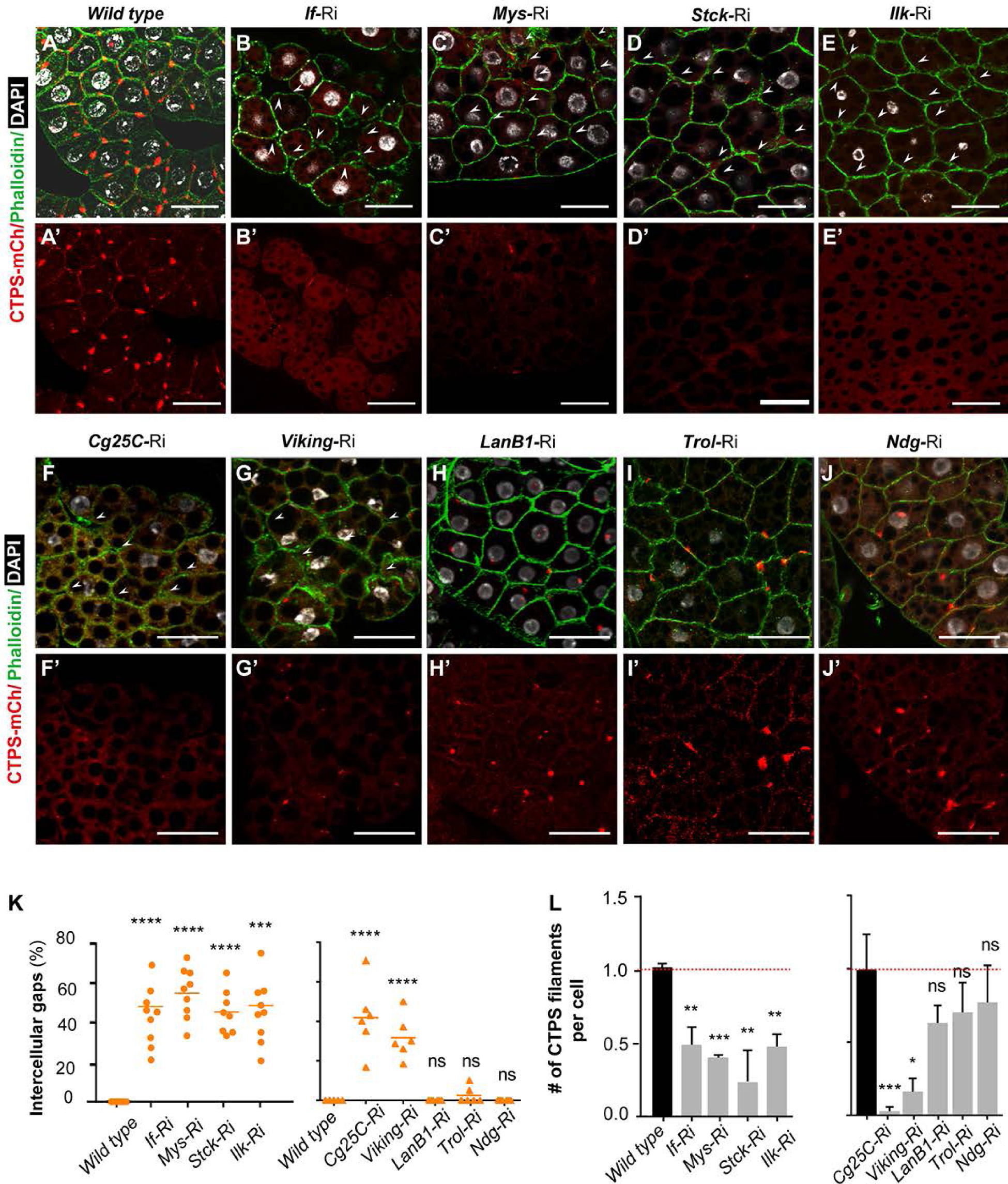
Integrin signaling promotes cytoophidium formation. (**A**-**J’**) Effect of reduced mRNA expression of some of the components in the Integrin signaling pathway on cytoophidium formation. Knock-down of mRNA expression was achieved via RNAi driven by the fat-body specific promoter *CgG4*. The fly lines examined were Integrin α2 (*If-*RNAi, **B**, **B’**), Integrin β (*Mys-*RNAi, **C**, **C’**), PINCH (*Stck-*RNAi, **D**, **D’**), ILK (*Ilk-*RNAi, **E**, **E’**), Collagen α1 (*Cg25C-*RNAi, **F**, **F’**), Collagen α2 (*Viking-*RNAi, **G**, **G’**), Laminin β (*LanB1-*RNAi, **H**, **H’**), Perlecan (*Trol-*RNAi, **I**, **I’**), Nidogen (*Ndg-*RNAi, **J**, **J’**). The *w^1118^* line was used as a wild-type control (**A**, **A’**). The *CTPS-*mCh line was crossed with these lines so that the effect on cytoophidium formation by the perturbations of these Integrin signaling components could be readily detected by visualizing mCherry fluorescence. Scale bars, 20 μm. (**K**) Quantifications of the adhesion defects from fat bodies at the third instar larval stage. The number of intercellular gaps per adipocyte was counted. Each point represents the number of gaps in a single fat body image. (L) Quantifications of the numbers of filaments per adipocyte of the fat bodies from the above lines. The value was normalized to the *CgG4*; *CTPS*-mCh>*w^1118^* line. All values are the means ± S.E.M. ns, no significance, * p < 0.05, ** p < 0.01, *** p < 0.001, **** p < 0.0001 by one-way ANOVA with a Tukey *post hoc* test.

Likewise, we also determined whether reduced expression of *Cg25C* (*Collagen at 25C*) or *Vkg* (*Viking*), which encodes the Collagen IV α1 chain or α2 chain, respectively, affected cytoophidium formation. In both cases, we observed defective adipocyte adhesion and concurrent absence of cytoophidia in fly lines with reduced expression of *Cg25C* or *Vkg* (Fig.6F, G, K, L). By contrast, we found that adipocyte adhesion was normal and cytoophidia were present in fly lines with reduced expression of Laminin, Perlecan, or Nidogen, all of which were also ECM components (Fig. 6H-L). The data thus suggest that Integrin signaling, and specifically that activated by Collagen IV rather than other ECM components, regulates cytoophidium formation.

### CTPS binds to integrin and colocalizes with integrin signaling complex

Next, we sought to determine how Integrin signaling promotes cytoophidium formation. To this end, we first examined whether a reduction in Integrin signaling may affect CTPS mRNA or protein expression level. Using qPCR, we found *CTPS* mRNA expression was not significantly altered in the fly lines expressing the *Mys-Ri*, *Stck-Ri*, or the *Ilk-Ri* allele (Fig. 7A). Likewise, Western-blotting analysis showed that CTPS protein levels were not significantly changed by reduced Integrin signaling in any of these fly lines (Fig. 7B, C). Together, these results show that Integrin signaling does not regulate CTPS at either the mRNA or protein expression level.

**Figure 7.**
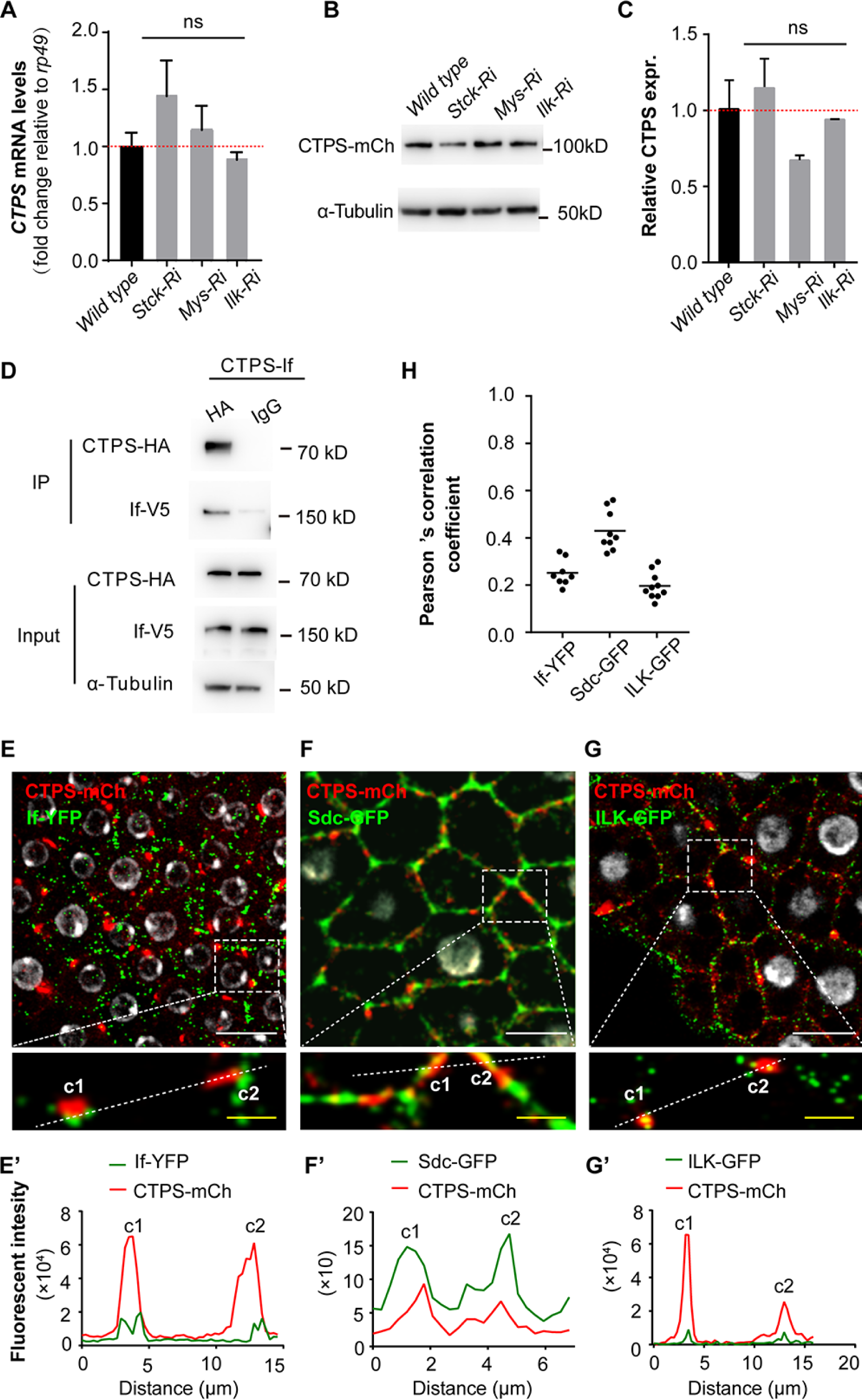
CTPS binds to integrin and colocalizes with integrin signaling complex. (**A**) Relative *CTPS* mRNA expression levels as measured by quantitative RT-PCR using fat bodies from the wild-type (*CgG4*; *CTPS*-mCh>*w^1118^*) and the fly lines with the indicated genotypes at the third instar stages (30 flies/genotype; n=3). Values were normalized to that from the wild type. (**B**, **C**) Relative CTPS protein expression levels as measured by Western Blotting analysis using fat bodies from the wild-type (*CgG4*; *CTPS*-mCh>*w^1118^*) and the fly lines with the indicated genotypes at the third instar stages (**B**, 30 flies/genotype; n=2). Values were normalized to that from the wild type. Anti-mCherry antibody was used to detect the expression of CTPS-mCherry fusion protein. Alpha-tubulin was used as an internal control. (**C**) Quantification of the CTPS protein relative expression levels. (**D**) Co-IP analysis of CTPS with associated proteins in S2 cells. S2 cells were transfected with the CTPS-HA-T2A-If-V5 plasmid. Antibodies against HA, V5 and α-tubulin were used for immunoblotting. (**E-G**) Fluorescent confocal imaging of adipocytes carrying the *CTPS-*mCh allele and one of the three alleles, including the If-YFP (**E**), or the Sdc-GFP (**F**), or the ILK-GFP (**G**). Areas in white dotted boxes were shown in close-up views to better illustrate colocalization between CTPS-mCH and If-YFP (**E’**), Sdc-GFP (**F’**), and lLK-GFP (**G’**). Note that labelled scans were performed along the white dotted lines in the close-up views to indicate individual cytoophidia adhering to each component of integrin adhesion complexes. (**H**) Quantification of co-localization analysis between CTPS and If-YFP, Sdc-GFP, and lLK-GFP. All values were the means ±S.E.M. ns, no significance by Student’s t-test. Scale bars, 20 μm (white), 5 μm (yellow).

Alternatively, Integrin signaling may promote cytoophidium formation by binding to its filament precursors and promote one or more steps during its polymerization. Therefore, we next tested whether CTPS could bind to Integrin. We “tagged” CTPS and Integrin with the HA and the V5 peptides, respectively, and expressed them in the S2 cells. These fusion proteins were then subjected to co-immunoprecipitation (Co-IP) assays to test their potential binding. We found that CTPS was able to bind to Integrin (Fig. 7D).

Moreover, if CTPS binds to Integrin proteins inside living cells, they should co-localize at the same places. To test this possibility, we crossed the If-GFP and the *CTPS-*mCh fly lines and examined Integrin and CTPS expression in the adipocytes. We found that Integrin and cytoophidia often colocalized in the fly adipocytes (Fig. 7E, E’, H). Likewise, cytoophidia also colocalized with other components of the Integrin signaling, including Syndecan (Sdc), an Integrin co-receptor, and ILK, which were labeled using the *Sdc*-GFP and the *Ilk*-GFP trap lines (Fig. 7F-H).

Taken all together, the above data show that Integrin signaling does not affect CTPS mRNA or protein expression; however, Integrin and most likely other components of the adhesion complex can bind to CTPS protein to promote filament formation.

## DISCUSSION

Tissue architecture is essential for organ physiology and function. How it is regulated by individual cell behavior, including cellular metabolism, has remained largely unclear. Here, we show that the metabolic enzyme CTPS regulates cell adhesion and tissue organization of the *Drosophila* adipose. CTPS forms filaments at the adipocyte cortex, corresponding to the Integrin-Collagen IV contact sites essential for adipocyte adhesion. Surprisingly, loss of CTPS expression causes defective cell adhesion and tissue dissociation, suggesting that CTPS promotes adipocyte adhesion. Interestingly, expression of mutant CTPS proteins, which have normal enzymatic activities but are unable to form filaments, also leads to defective adipocyte adhesion, suggesting that the polymer form, rather than the absolute amount, of CTPS proteins, is essential for adhesion and tissue architecture. We show that cytoophidia promote Collagen IV mRNA expression and thus its deposition to strengthen Integrin-Collagen IV adhesion. Moreover, a reduction of Integrin signaling due to loss of Integrin or other components of the signaling pathway leads to failure of cytoophidium formation. We show that Integrin signaling does not affect CTPS mRNA expression, but increases the protein expression by promoting its filament formation, which stabilizes the otherwise labile monomers and tetramers. This is supported by the binding and colocalization of components of the Integrin adhesion complex with cytoophidium.

### Coupling Adipose Tissue Architecture and Metabolism via Integrin Feedback Signaling

One of our most striking observations was the cortical location of cytoophidia, which were aligned with adipocyte-matrix contact sites known to require Integrin-Collagen IV interactions (Dai et al. 2017). Indeed, loss of Integrin signaling activities, for example, as a result from reduced expression of Collagen IV, Integrin, Syndecan, or downstream signaling components PINCH, or ILK, all led to a failure of cytoophidium formation. These data thus conclusively demonstrate that Integrin-Collagen IV signaling promotes cytoophidium formation (Fig. 8).

**Figure 8.**
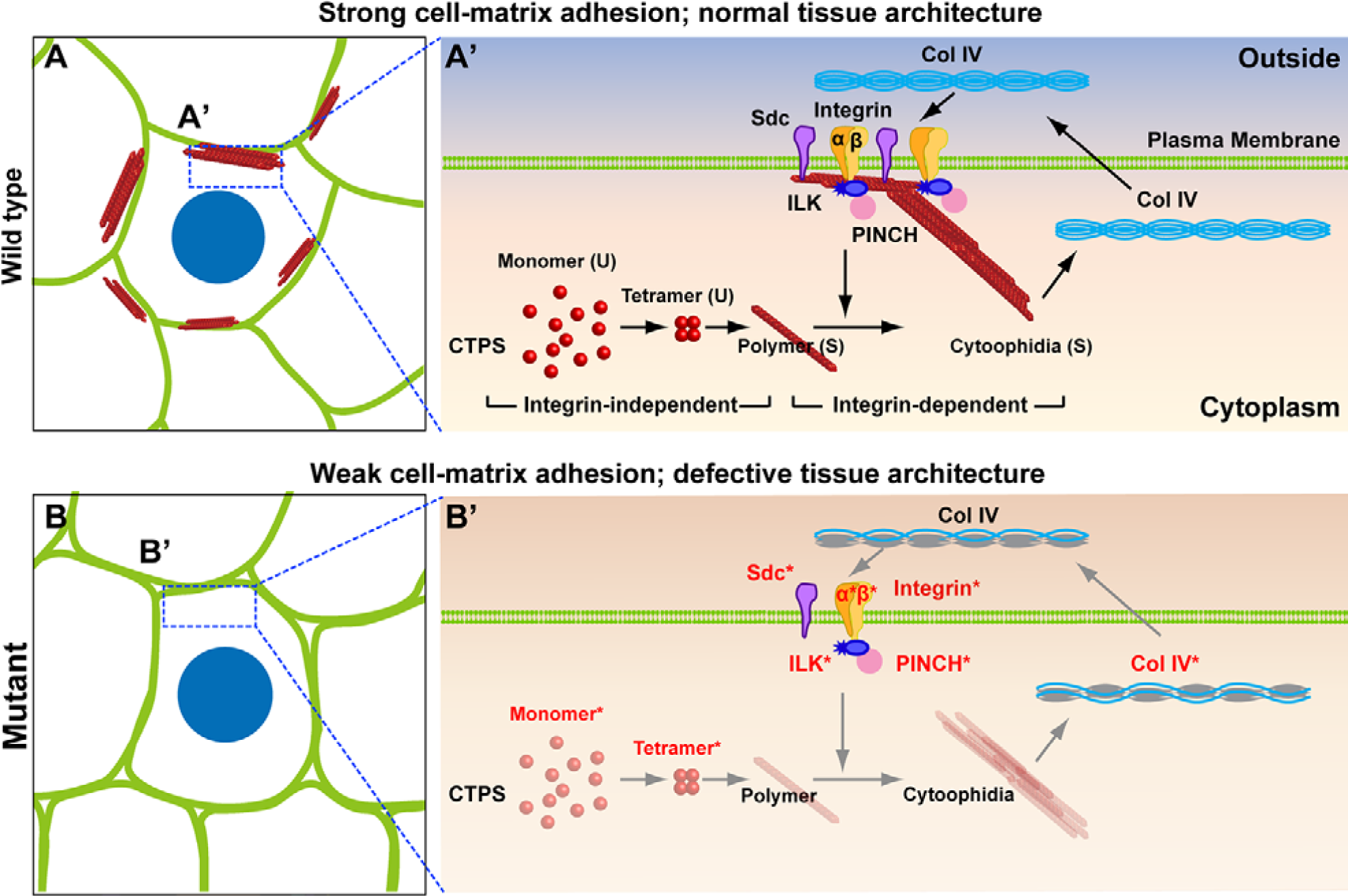
Working model of the CTPS-Integrin signaling feedback loop that regulates adipocyte adhesion in fly fat-body. (**A**) A schematic diagram showing the control adipose tissue, whose organization and architecture is based on tight adipocyte-matrix adhesions as mediated by Collagen IV-Integrin binding. (**A’**) A signaling feedback looping involving both cytoophidia and Collagen IV in the extracellular matrix. cytoophidium promotes Collagen IV mRNA expression and protein deposition, which in turn, via binding to Integrin, is essential for adipocyte adhesion. Binding by Collagen IV activates Integrin signaling, which via downstream components including PINCH and ILK, promotes cytoophidium formation. Furthermore, cytoophidium formation is a multistep process, including an early, Integrin-independent step where unstable (U) monomers and tetramers form stable (S) polymers. It also include a late step that is Integrin signaling dependent, where the polymer forms of CTPS undergo a higher order assembling step and form the microscopically more visible filaments. (**B**) Mutant adipose tissue with weakened cell-matrix adhesion and defective organization and tissue architecture. (**B’**) Adipocyte adhesion can be weakened by any one of the steps along the signaling feedback loop. The steps that were experimentally manipulated in the current study were marked in red and by an asterisk (*). As a result, Integrin-mediated cell adhesion is reduced, resulting in defective adipose architecture.

Remarkably, the formation of cytoophidia also promotes Integrin-Collagen IV-mediated adipocyte adhesion. In the absence of cytoophidia, whether due to reduced CTPS expression or forceful expression of a mutant CTPS protein that prevents the polymerization event, in turn, led to reduced Integrin signaling, weakened adipocyte adhesion, and defective adipose tissue architecture (Fig. 8). Together, our data demonstrate that there exists a signaling feedback loop between cytoophidium formation and Collagen IV in the matrix, by which intrinsic cellular metabolism of adipocytes is coupled to extrinsic tissue architecture of the adipose (Fig. 8). We speculate that, by coupling these two essential characteristics, the fly adipose, and possibly other tissue and organ systems can best match their structure and physiological function to operate at an optimal condition.

Our data show that the amount of cytoophidia correlated with Collagen IV protein expression levels, suggesting that cytoophidia promote Collagen IV expression. Interestingly, a similar, though weaker, correlation was also observed between cytoophidium amount and Integrin expression levels (Fig. 5). This difference in the amount of promotion between Collagen IV and Integrin protein expression could be partly explained by the notion that Collagen IV functions upstream of Integrin and thus its increase due to cytoophidium formation secondarily upregulated Integrin expression levels, although to a lesser extent.

It remains unclear the mechanism by which cytoophidia regulate Integrin signaling in fly adipocytes. Surprisingly, CTPS mRNA levels, and possibly the amount of cytoophidia, appeared to be directly correlated with Collagen IV mRNA expression levels (Fig. 5K). These data suggest that CTPS promote Collagen IV expression at the transcriptional level, presumably indirectly via other protein partners. We await with great anticipation the exciting discoveries of these protein partners that, together with CTPS, may regulate Collagen IV expression and Integrin signaling.

### Integrin signaling-independent and dependent stages during cytoophidium formation

We show that the replacement of the wild-type CTPS protein by a mutant form, in which H^355^ is converted into A^355^ or R^355^, in the *CTPS*^MU^-mCh or the *CgG4*/CTPS^MU-OE^ fly lines caused a failure of filament formation in the fly adipocytes. The data are thus consistent with a previous report showing that the H^355^ is a crucial amino-acid residue for tetramer formation in human CTPS proteins (Lynch et al. 2017). Moreover, CTPS protein levels were also reduced in the *CTPS*^MU^-mCh adipocytes when compared to normal. Interestingly, despite a reduction in protein expression, CTPS mRNA expression in the *CTPS*^MU^-mCh adipocytes was similar to that in the control (Fig. 5K). These data thus suggest that filament formation promotes CTPS protein expression at a post-transcriptional level, presumably by stabilizing the highly labile monomers and tetramers (Fig. 8).

Remarkably, however, despite a positive control of Integrin signaling on cytoophidium formation, we observed no reduction in CTPS mRNA or protein expression levels in the fly lines, in which Integrin signaling was downregulated. Together, these data suggest that cytoophidium formation is a multi-step process. Specifically, it includes an early, Integrin-independent step whereby the unstable monomers and tetramers form the stable polymers; and a late, Integrin-dependent step whereby polymers further bundle into a higher-order filament, both of which are stable (Fig. 8). Moreover, considering that the endogenous, wild-type CTPS proteins would have also been expressed by *CgG4*/CTPS^MU-OE^ adipocytes, we concluded that the mutant form of CTPS must have functioned as a dominant-negative and prevented filament formation of the wild-type CTPS proteins.

At present, it remains unclear how Integrin signaling may promote cytoophidium formation. It is possible that Integrin regulation may be direct or indirect as a result of interactions with the proto-oncogene Myc, the ubiquitin E3 ligase (Cbl), and activated Cdc42-associated kinase (Ack), all of which are known to regulate cytoophidium formation in various settings(Azzam and Liu 2013; Strochlic et al. 2014; Pai et al. 2016). Alternatively, Integrin signaling may regulate cytoophidium formation at a metabolic level. Indeed, Integrin signaling is known to regulate glucose metabolism and glutamine transportation, whose substrates may be used by CTPS in the adipocytes (Low and Taylor 1998; Williams et al. 2015; Kim et al. 2018; Fu et al. 2019; Bae et al. 2020; Zhou et al. 2021). These are interesting possibilities that we anticipate to be scrutinized by future studies.

## MATERIALS AND METHODS

### Generation of transgenic flies

The CRISPR/Cas9 technology was used to establish both the C-terminal mCherry-4V5 tagged CTPS knock-in fly and C-terminal mCherry-3HA tagged CTPS H355A mutation knock-in fly according to homology-directed repair procedures as previously described (Bassett et al. 2013) at Fungene Biotech (http://www.fgbiotech.com). For C-terminal mChe-4V5 tagged CTPS knock-in fly generation, the two single strain guide RNA (sgRNA) were designed with CRISPR Optimal Target Finder (Gratz et al. 2014). sgRNA1: GCCATAAGTAAACTAGTGAACGG, sgRNA2: CCTAAAGTGTTTAACATCCGATT. For donor vector construction, the mCh-4V5 cassette amplified from pBluescript SK-mCherry-4V5 (from Fungene Biotech, unpublished) was cloned into pBSK (-) vector containing CTPS arm regions. The donor vector pBSK CTPS-mCherry-4V5, sgRNAs, and Cas 9 mRNA were injected into W1118 embryos. PCR sequencing was performed to validate if the offspring flies carried mCherry-4V5 insertion. Three steps were carried out to generate C-terminal mCherry-3HA tagged CTPS H355A mutation knock-in fly. The first step, CTPS-KO-attp fly were constructed. The four single strain guide RNA (sgRNA) were designed with CRISPR Optimal Target Finder. CTPS attp-RFP 5sgRNA1: CGATGGCGCCGAGGTGGATCTGG, CTPS attp-RFP 5sgRNA2 TGGATCTGGGAAACTATGAACGG, CTPS attp-RFP 3sgRNA1: GCGGGCAGCAAGAACGGTATTGG, CTPS attp-RFP 3sgRNA2: ACGGTATTGGAAATAGTGCGTGG. For donor vector construction, the attP-RFP-Loxp cassette amplified from the plasmid attP-RFP (from Fungene Biotech, unpublished) with primer attp-RFP-F and Loxp-R was cloned into pBSK (-) vector containing CTPS KO arm regions. The donor vector pBSK CTPS-KO-attp, sgRNAs and cas 9 mRNA were injected into *w^1118^* embryos. PCR sequencing was performed to validate if the offspring flies carried attp-RFP-Loxp insertion. Step 2, the cDNAs encoding Drosophila CTPS was produced by RT-PCR using the total RNAs extracted from the *w^1118^* line. Then, mCherry-3HA sequence was ligated to the C-terminal of CTPS. H355A point mutation was introduced into pUAST-attB-CTPS plasmid. Finally, site-specific integration of pUASTattB-CTPSH355A plasmid into CTPS-KO-attp germ line by co-injection with phiC31integrase RNA. Step 3, the offspring flies from step2 were crossed with cre fly to remove non-essential sequence between two Loxp. Thus, C-terminal mCherry-3HA tagged CTPS H355A mutation knock-in fly were finally generated.

To generate transgenic UAS-CTPSWT and UAS-CTPS^MU^ flies, the cDNAs encoding Drosophila CTPS was produced by RT-PCR using the total RNAs extracted from the *w^1118^* line (#3605, from the Bloomington Drosophila Stock Center). The oligonucleotide primers used were as follows: CTPS sense 5’-ttcgttaacagatctgcggccgcatgaaatacatcctggtaact-3’, antisense 5’-ttcacaaagatcctctagaggtacccttgtacagctcgtccatgc-3’. The PCR products were digested with NotI and KpnI and cloned into the pUASTattB vector for the expression of mCherry-HA-tagged CTPS protein. H355R point mutation was introduced into pUAST-attB-CTPS plasmid by using Mut Express II Fast Mutagenesis Kit V2 (Vazyme) following the manufacturer’s instructions. The mutagenic oligonucleotide primers used was sense 5’GAGCAAGTACCGGAAGGAGTGGCAGAAGCTATGCGATAGCCAT-3’, antisense 5’-TGCCACTCCTTCCGGTACTTGCTCGGCTCAGAATGCAAAGTTT-3’.

Then, site-specific integration of pUASTattB-CTPS or pUASTattB-CTPS^H355-R^ plasmid into fly germ line (attp2) that contain attp lading sits was carried out by co-injection with phiC31integrase RNA as previously described 48 at the Core Facility of *Drosophila* Resource and Technology, SIBCB, CAS.

### Fly strains

The GAL4/UAS system (Brand and Perrimon 1993) was use for adipocyte-specific expression or RNAi knockdown of the desired genes. The *Cg*-*GAL4* driver lines were obtained from the Bloomington *Drosophila* Stock Center (BDSC; Department of Biology, Indiana University, Bloomington, IN). The *Cg*-*GAL4* line was crossed with the *w^1118^* flies to generate the *Cg*-*GAL4*> *w^1118^* control lines. The fly lines were obtained from the Bloomington *Drosophila* Stock Center, including UAS-*CTPS*-RNAi (stock number 31924), UAS-if-RNAi (stock number 27544) UAS-*mys*-RNAi (stock number 27735) and tGPH (stock number 8163), or from the Vienna Drosophila RNAi Center, including UAS*-ilk*-RNAi (V35374). UAS*-mys*-RNAi*^VDRC.v29619^*, UAS-*stck*-_RNAi_*TRiP.JF01096*_, UAS*-ilk*-RNAi_*TRiP.GL00288*_, UAS-*Vkg*-RNAi_*NIG.16885R-3*_, UAS-_ *Cg25C*-RNAi^VDRC.v28369^, UAS-*LanB1*-RNAi^VDRC.v23121^, UAS-*Ndg*-_RNAi_Trip.HMJ24142 _UAS-*Trol*-RNAi_VDRC.v24549_, Vkg_G454_.GFP, Ndg.sGFP_ fTRG.638, If^CPTI-004152.YFP^, and ILK^ZCL3192.GFP^ were kind gifts from José C. Pastor-Pareja at Tsinghua University.

### Fly husbandry and diet preparation

Fly lines were raised on the standard yeast-cornmeal-agar food. To make sure larvae used in this study at the desired developmental stage, we restricted egg collections by allowing female to lay eggs for less 4 hours. All flies were cultured at 25°C with 50% humidity under a 12 h/12 h light/dark cycle.

### Protein expression constructs

For expression of the wild-type CTPS, and If, cDNA fragments were amplified by RT-PCR using the total RNAs extracted from the ***w^1118^*** line. The oligonucleotide primers used were as follows: *CTPS* sense 5’-ttcgttaacagatctgcggccgcatgaaatacatcctggtaact-3’, antisense 5’-ttcacaaagatcctctagaggtacccttgtacagctcgtccatgc-3’. *If* sense 5’-ggagaatcccggccctgcgatgagtggagattccatccaccg-3’, antisense 5’-gggataggcttaccttcgaacaggtgctcgtcgccgtg-3’. *CTPS* cDNA was cloned into pAc5.1 plasmid, then the HA and T2A sequences were added in the 3’ end of *CTPS* followed by inserting *If*-V5 fragments.

### Immunohistochemistry

The fat body from the 2^nd^ or 3^rd^ instar larvae were dissected in Grace’s Medium and then fixed in 4% formaldehyde in PBS for 15 min before immunofluorescence staining. For membrane staining, fixed fat bodies were washed twice for 5 min in PBS and then were incubated with 0.165 μM Alexa Fluor 488 phalloidin or Alexa Fluor 633 phalloidin (Invitrogen) in PBSTG for 30 min at RT. Then samples were rinsed in PBST twice for 5min each and mounted in Vecta shield with DAPI (Invitrogen). For cytoophida immunostaining, goat anti CTPS (1:400, Santa Cruz Catalogue no.33304) was used as the primary antibody, and Cy5-AffiniPure donkey anti-goat IgG (1:1000, Jackson Immuno Research Laboratories, Inc., Catalogue no. 705-175-147) as the secondary antibody. Tissues were mounted in the anti-fading mounting medium with DAPI (Invitrogen). Then samples were rinsed in PBS twice for 5min each and mounted in Vecta shield with DAPI (Vector Labs).

### S2 cell culture and transfection

*Drosophila* S2 cells were cultured in Schneider’s medium (Invitrogen) containing 10% heat-inactivated fetal bovine serum (Invitrogen) at 25 °C. Cell transfections were carried out by using FuGENE HD Transfection Reagent (E2311, Promega, WI) according to the manufacturer’s instructions.

### Co-immunoprecipitation

S2 Cells were harvested 24 h after transfection with pAC 5.1 CTPS-HA-T2A-If-V5. Then cells were lysed in prechilled Co-IP lysis buffer (20mM Tris-HCl, pH 7.4, 50mM NaCl, 2mM MgCl2, 1% [vol/vol] NP40, 0.5% [mass/vol] sodium deoxycholate, and 0.1% [mass/vol] sodium dodecyl sulfate) containing 1X protease inhibitor cocktail (Bimake) for 2 hours on ice. After centrifuging 15000g at 4oC, the supernatants were collected and incubated with anti HA magnetic beads (Bimake) or IgG bound protein A/G magnetic beads (Bimake) followed by an overnight rotation at 4oC. Then the beads were washed three times with 1 ml PBST (0.5% Tween20 in PBS) for 5 minutes, followed by SDS-PAGE and immunoblotting analysis after elution by boiling in 50 μl 1× SDS loading buffer.

### Imaging and image analysis

Fluorescent images were obtained by confocal laser-scanning microscopy (Leica SP8). Super resolution images were obtained by TCS SP8 STED 3X super resolution stimulated emission depletion microscopy. For quantification of intercellular gaps, images were acquired at 40x oil object and used to score the percentage of gaps. Each data point represents the percentage of gaps analyzed from one image. For quantification of CTPS-mCherry filaments, fluorescence intensity was measured in 40X confocal images. FIJI-ImageJ was used to measure signal intensity in an ROI containing CTPS-mCherry filaments in one image. Fluorescence intensity was normalized by ROI area in each image. For quantification of abundance of CTPS-mCherry filaments, the number of CTPS mCherry-containing filaments was counted in cells of 40X confocal images with FIJI-ImageJ. The data represents filaments number in one cell normalized by cell number in one such image. The calculation of Pearson’s correlation coefficients after mannual Costes thresholding were based on intensity/voxel with Imaris. Each data point represents one image.

### Western blot

Larvae or fat body tissues were extracted in RIPA buffer (150 mM NaCl, 1% NP-40, 0.5% sodium deoxycholate, 0.1% SDS, 50 mM Tris-HCl, pH 7.4) using a Tissuelyser-24 grinder (Jingxin, Shanghai, China). After centrifugation at 15,000g at 4°C for 20 min, the supernatants were subjected to separation by SDS-PAGE before immunoblotting analysis. The following antibodies were used for immunoblotting: primary antibodies included mouse anti-V5 antibody (1:1000, Invitrogen, Catalogue no. 1461501), mouse anti-α-Tubulin antibody (1:1000; Sigma, Catalogue no. T6199), mouse anti-HA (1:1000, Santa Cruz Biotech., Catalogue no. sc-7392), mouse anti-mCherry (1:1000, Abbkine Scientific Co., Ltd Catalogue no. A02080), V5-Tag (D3H8Q) mAb Rabbit (1:2000, CST, Catalogue no.13202S). Secondary antibodies include Horseradish peroxidase (HRP)-conjugated anti-mouse IgG, (1:2000, Cell Signaling, Catalogue no.7076), Nonsaturated bands were quantified on ImageJ (National Institutes of Health) and presented as a ratio in relation to the internal reference α-Tubulin. At least two-three biological replicates were quantified.

### Quantitative RT-PCR

Total RNAs were prepared from the lysate of larval fat body tissues using the TRIzol reagent (TransGen Biotech, Beijing, China). cDNAs were synthesized with PrimeScript RT Master mix (Takara) followed by adding template RNA. 2X SYBR Green PCR Master Mix was purchased from Bimake. Real-time quantitative PCR was conducted using the QuantStudion^TM^ 7 flex System (Applied Biosytstems). For normalization, *rp49* was utilized as the internal control. The oligonucleotide primers used were as follows:

*rp49*: sense 5’-TCCTACCAGCTTCAAGATGACC-3’, antisense 5’-CACGTTGTGCACCAGGAACT-3’;

*CTPS*: sense 5’-GAGTGATTGCCTCCTCGTTC-3’, antisense 5’-TCCAAAAACCGTTCATAGTT-3’.

*Viking*: sense 5’-GGAGATGTCGGCGAGTATGG-3’, antisense 5’-GAACTCCGTTCTFFCCAGGT-3’.

*Cg25C*: sense 5’-GGATCCATCGGACCCATTGG-3’, antisense 5’-GTCCAFFAGCACCAGCGG-3’.

### 3D reconstruction

The model tetramer structure of *Drosophila* CTPS is built by Prime of Schrödinger Suite 2018-4, with human CTPS structure (PDB code: 5U03) as the template. The octamer structure of *Drosophila* CTPS was obtained by mapping the *Drosophila* CTPS model into the EM density map of human cytoophidium (EMD-8474) via “Fit in map” tool of UCSF Chimera. The resulting *Drosophila* CTPS octamer model was further processed by “Protein preparation wizard” in Schrödinger Suite, to obtain a proper hydrogen bond network and to perform an energy optimization.

### Statistical analysis

All data are presented as the mean ± standard errors of the mean (s.e.m) from at least three independent experiments. Statistical analysis between each genotype and the controls was determined by unpaired two-tailed Student’s t-test, whereas multiple comparisons between genotypes were determined by one-way or two way ANOVA with a Tukey post hoc test in GraphPad Prism 7.0. P<0.05 was considered to be statistically significant.

## ACKNOWLEDGEMENTS

We thank Shuang Zhou, Xiaoming Li, and Rui Wang for technical assistance. We also thank the Core Imaging Facility at the National Center for Protein Science Shanghai (NCPSS) and the Molecular Imaging Core Facility (MICF) at School of Life Science and Technology, ShanghaiTech University for providing technical support. We thank Nolan Liu for proof-reading. We are grateful for comments from Sibao Wang and Tiffany Horng.

## AUTHOR CONTRIBUTIONS

J.L. and J.L.L. conceived the studies. J.L. designed the studies and performed most of the experiments. Y. Zhang performed biochemical experiments including qRT-PCR, Western Blot and Co-IP experiment. Y. Zhou and Q.W. constructed plasmids for transgenic fly lines. K.D. and S.Z. built 3D reconstruction of CTPS. J.L. drafted the manuscript. P.L. and J.L.L. revised the manuscript.

## DATA AND MATERIALS AVAILABILITY

All data are provided in the text or supplementary materials.

## SUPPLEMENTARY INFORMATION

### SUPPLEMENTARY FIGURES S1-S4

**Supplemental Figure S1.**
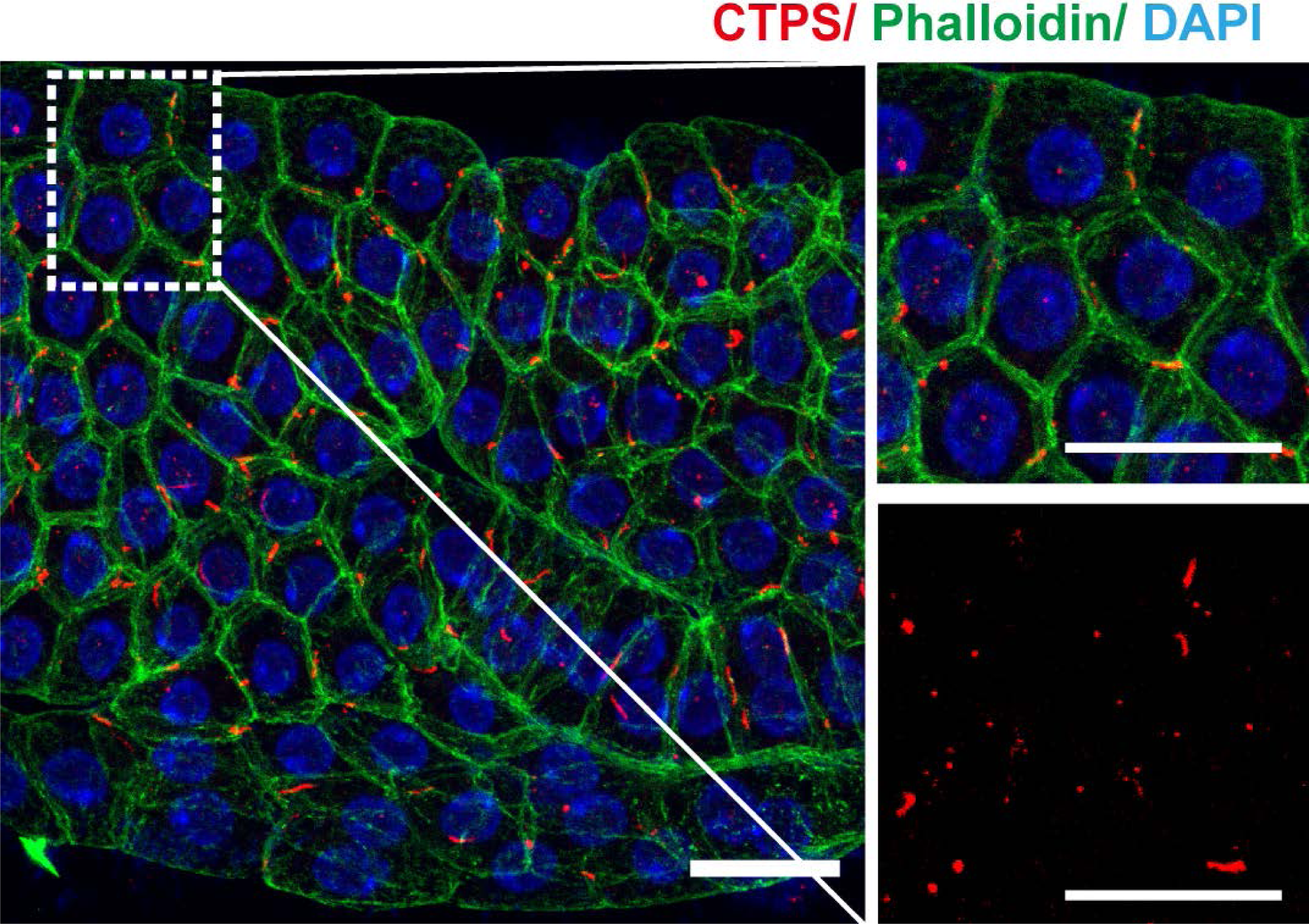
Cytoophidia locate intercellular contacts in *Drosophila* adipocytes. Confocal images of the fat bodies of the second instar larvae of *w^1118^* show the localization of cytoophidia adjacent to the plasma membrane of the fat body cells. cytoophidia are stained with an anti-CTPS antibody (red). The plasma membrane of the cells is stained with phalloidin (green). Nuclei are stained with DAPI (blue). The area marked by the white square is magnified in the right panel. Scale bars, 20 μm.

**Supplemental Figure S2.**
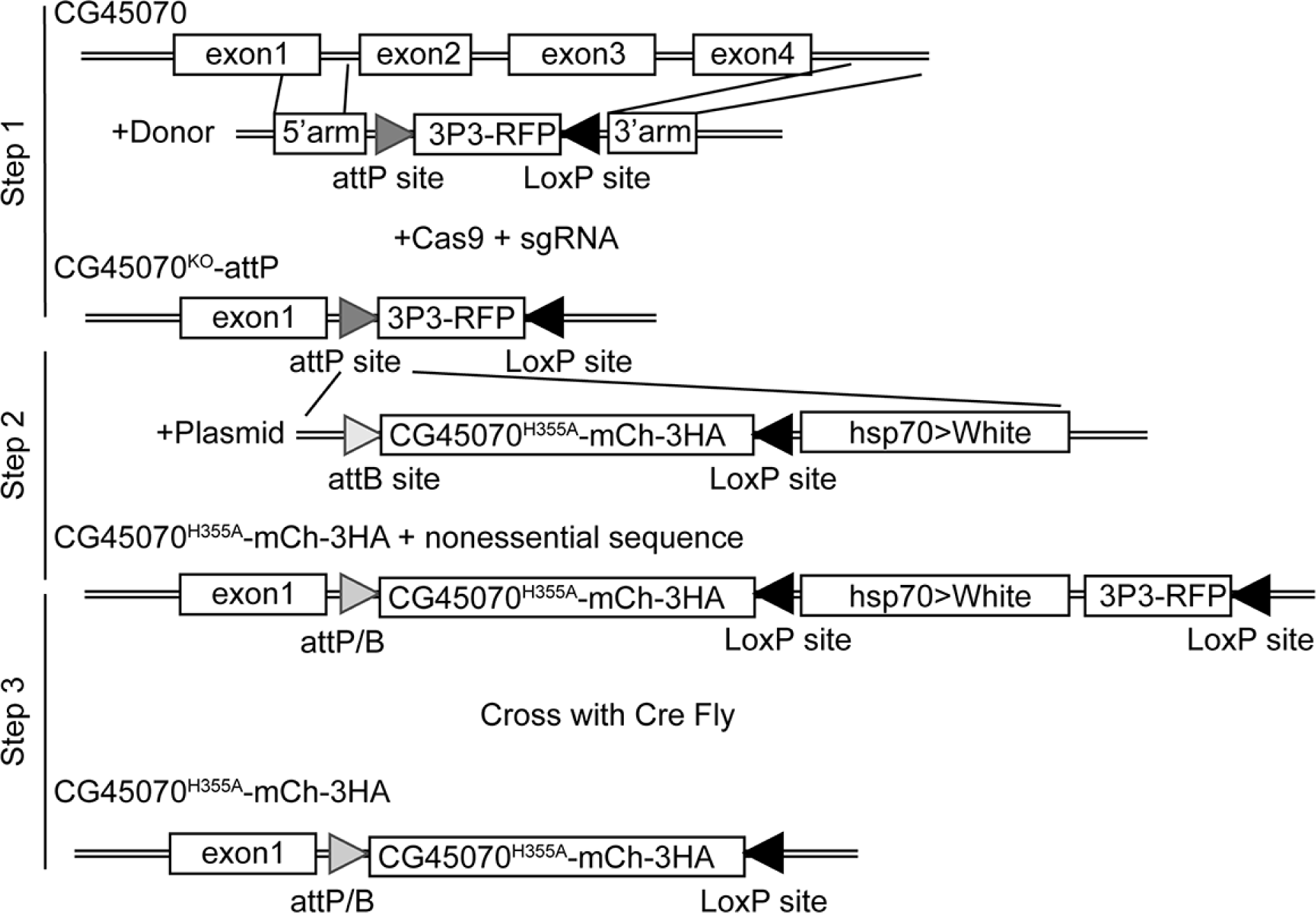
Construction of H355A mutation CTPS knock-in fly. Schematic of CRISPR/Cas-mediated CTPS-H355A-mCherry-3×HA (*CTPS*^MU^-mCh) generation. Step1: Generation of CTPS (CG45070) knock out fly line via CRISPR/Cas technology. Step2: cDNA with H355A mutation tagged by mCherry and 3HA was cloned into vector pUASTattB containing attB site. Site-specific integration of the resulting construct into attP40 flies at attP landing site was carried out by coinjection with phiC31-integrase RNA. Step3: Removal of non-essential sequence by Cre/Loxp system to generate *CTPS*^MU^-mCh fly.

**Supplemental Figure S3:**
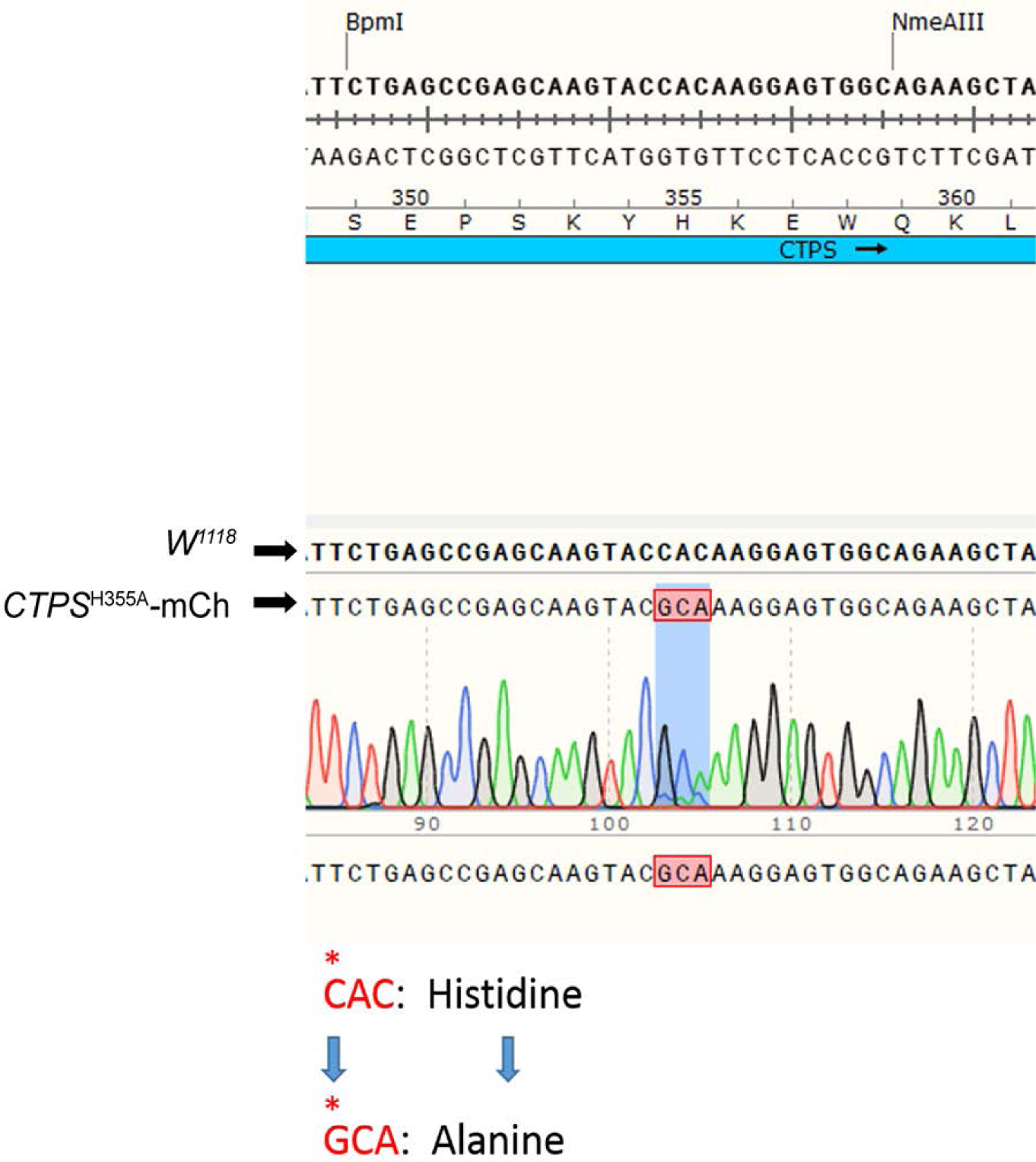
Sequencing analysis. Sequencing analysis of genomic DNA to confirm CTPS H355A point mutation in *CTPS*^MU^-mCh fly.

**Supplemental Figure S4:**
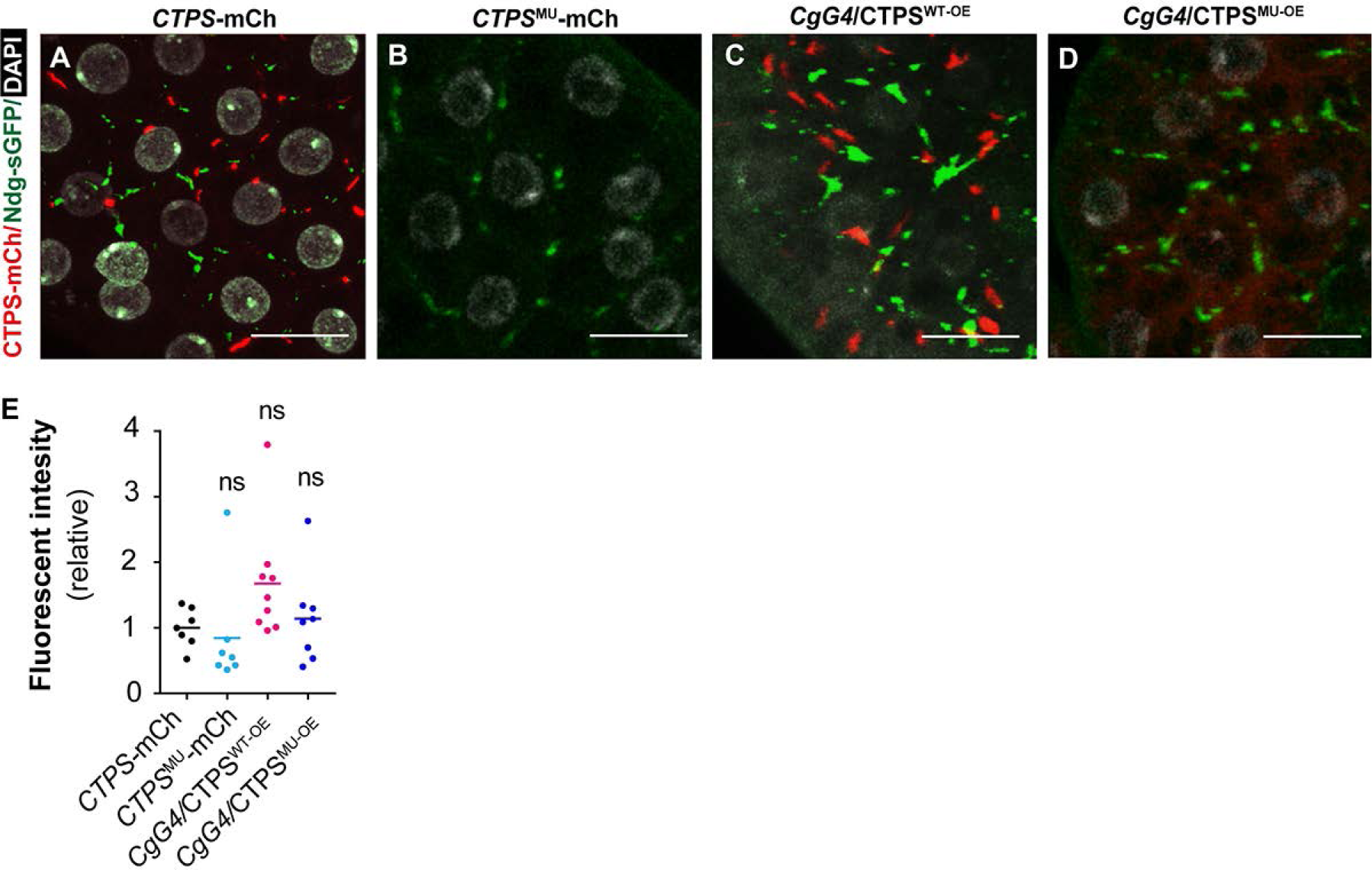
Effects of CTPS on Nidogen distribution. (**A**-**D**) Representative confocal images of fat bodies from the 3rd instar larvae showing the intercellular presence of Ndg-sGFP (green) upon knock in of mutant CTPS, overexpression of wild type CTPS, or mutant CTPS by *Cg GAL4*. Scale bar, 20 µm. (**E**) Quantitative analysis of Ndg-sGFP intensity from A-D upon knock-in of mutant CTPS, overexpression of wild-type CTPS, or mutant CTPS. The value was normalized to the *CTPS*-mCh line. All values are the means ± S.E.M. ns, no significance by one-way ANOVA with a Tukey post hoc test.

